# CasGen: A Regularized Generative Model for CRISPR Cas Protein Design with Classification and Margin-Based Optimization

**DOI:** 10.1101/2025.02.28.640911

**Authors:** Bharani Nammi, Vindi M. Jayasinghe-Arachchige, Sita Sirisha Madugula, Maria Artiles, Charlene Norgan Radler, Tyler Pham, Jin Liu, Shouyi Wang

## Abstract

Clustered Regularly Interspaced Short Palindromic Repeats (CRISPR)-associated proteins (Cas) systems have revolutionized genome editing by providing high precision and versatility. However, most genome editing applications rely on a limited number of well-characterized Cas9 and Cas12 variants, constraining the potential for broader genome engineering applications. In this study, we extensively explored Cas9 and Cas12 proteins and developed CasGen, a novel transformer-based deep generative model with margin-based latent space regularization to enhance the quality of newly generative Cas9 and Cas12 proteins. Specifically, CasGen employs a strategies that combine classification to filter out non-Cas sequences, Bayesian optimization of the latent space to guide functionally relevant designs, and thorough structural validation using AlphaFold-based analyses to ensure robust protein generation. We collected a comprehensive dataset with 3,021 Cas9, 597 Cas12, and 597 Non-Cas protein sequences from reputable biological databases such as InterPro and PDB. To validate the generated proteins, we performed sequence alignment using the BLAST tool to ensure novelty and filter out highly similar sequences to existing Cas proteins. Structural prediction using AlphaFold2 and AlphaFold3 confirmed that the generated proteins exhibit high structural similarity to known Cas9 and Cas12 variants, with TM-scores between 0.70 and 0.85 and root-mean-square deviation (RMSD) values below 2.00 Å. Sequence identity analysis further demonstrated that the generated Cas9 orthologs exhibited 28% to 55% identity with known variants, while Cas12a variants show up to 48% identity. Our results demonstrate that the proposed Cas generative model has significant potential to expand the genome editing toolkit by designing diverse Cas proteins that retain functional integrity. The developed deep generative approach offers a promising avenue for synthetic biology and therapeutic applications, enableling the development of more precise and versatile Cas-based genome editing tools.

## 1 Introduction

Clustered Regularly Interspaced Short Palindromic Repeats (CRISPR)-associated protein 9 (Cas9) systems, which originate from adaptive immune mechanisms in bacteria and archaea, have revolutionized genome editing due to their precision and versatility^1–3^. Comprehensive genomic analyzes have significantly expanded the understanding of Cas9 diversity, identifying Cas9 orthologs in 653 bacterial strains in 347 species^1^. More recent studies have further demonstrated this diversity, revealing Cas9 genes in 92,140 loci from over a million microbial genomes, which have been further classified into 2,779 distinct groups according to sequence identity, with functional Protospacer Adjacent Motifs (PAMs) predicted for 2,546 groups^3^. These findings highlight the vast potential of a broader spectrum of Cas9 proteins, each with unique targeting capabilities, which could be optimized for specialized genome-editing applications.

Despite the diversity of Cas9 enzymes identified across various bacterial and archaeal species, current genome-editing technologies rely on only a limited set of well-characterized Cas variants. Among them, *Streptococcus pyogenes* Cas9 (SpCas9) is the most widely used due to its high editing efficiency and ease of use across experimental models^1,2,4–7^. Each Cas9 variant has its own set of limitations, some of which are shared across systems while others are unique to specific variants. For instance, SpCas9 and other commonly used Cas9 proteins face limitations, including off-target effects, strict PAM sequence dependence, and large protein size, which can hinder efficient delivery into cells^8–14^. These limitations have driven researchers to pursue two complementary strategies: engineering existing Cas proteins to enhance their functionality and discovering new Cas variants with improved properties. Advancements in genomic sequencing and bioinformatics have made it possible to explore less-studied Cas9 orthologs, which may possess unique enzymatic properties suited for specific genome-editing tasks. Rational protein design, directed evolution, and computational modeling have all been employed to optimize Cas9 enzymes, yet the search for functionally superior variants remains a significant challenge^1–5,15–17^. While these approaches have led to enhanced Cas enzymes with modified PAM preferences, improved specificity, and reduced off-target activity, they remain limited by the available sequence diversity and experimental screening bottlenecks.

To overcome these challenges, machine learning and deep generative models have emerged as powerful tools for exploring the vast sequence space of Cas proteins, enabling the design of novel variants with desirable properties. Advancements in deep learning have enabled computational models to predict protein structure, function, and evolutionary relationships with unprecedented accuracy. Generative models, including Variational Autoencoders (VAEs), Generative Adversarial Networks (GANs), and diffusion models, have been successfully applied to de novo protein design, capturing complex sequence-function relationships that are difficult to model through traditional bioinformatics techniques. Large-scale protein language models such as ProtGPT and ESM-1b have further demonstrated the ability to generate biologically meaningful sequences with functional attributes^18–21^. Several proof-of-concept studies have also validated AI-generated sequences experimentally, confirming that computational methods can yield enzymes with promising characteristics for genome engineering^22–24^. However, existing models are generally trained for general protein data generation and lack of dedicated training and learning of Cas protein to design high quality complex Cas proteins. Harnessing the cutting-edge generative models specifically for CRISPR-associated proteins opens new avenues for optimizing traits such as nuclease specificity, PAM compatibility, and delivery efficiency.

In our previous study, we conducted an extensive study on CRISPR-Cas protein pattern recognition and successfully developed a transformer-encoder-based deep learning model capable of identifying Cas protein sequences with high accuracy, exceeding 98%^25^. In this study, we introduce CasGen, a novel Transformer-based deep generative model for designing Cas9 and Cas12 variants. Our model integrates margin-based latent space regularization to enhance the quality of generated protein sequences while ensuring they retain essential functional characteristics. Unlike conventional generative approaches that prioritize sequence diversity without explicitly enforcing structural constraints, CasGen incorporates a classification network to distinguish Cas-like from non-Cas sequences, followed by Bayesian optimization to refine the latent space, guiding sequence generation toward functionally relevant proteins. Unlike existing general protein generative tools, our framework is tailored specifically for Cas protein engineering. Our study presents a systematic approach to expanding the genome editing toolkit through deep generative modeling. The key contributions of this work include: 1) a novel Transformer-based generative model (CasGen) with margin-based latent space regularization for Cas protein design; 2) a classification-enhanced generative pipeline that filters non-Cas sequences to improve sequence specificity; 3) Bayesian optimization of the latent space, ensuring generated sequences are structurally viable and functionally relevant; 4) extensive computational validation using AlphaFold-based structural modeling and sequence alignment techniques^26,27^. By integrating generative AI with structural validation, CasGen represents a promising step toward designing novel Cas proteins with enhanced specificity, delivery potential, and genome-editing capabilities. This approach opens new avenues for synthetic biology and therapeutic genome editing, enabling the rapid development of next-generation CRISPR tools.

## 2 Related Works

The field of protein sequence generation and design has seen rapid advancements, driven by the evolution of machine learning (ML) and deep learning (DL) models. This section reviews the key generative approaches and contextualizes our work within this growing landscape.

Traditional protein sequence generation relied on multiple sequence alignments (MSAs) and statistical modeling. Rao et al. (2021) introduced the MSA Transformer, a pioneering deep learning model that leverages MSAs to generate protein sequences by utilizing an masked language modeling (MLM) objective^28^. This approach effectively captures co-evolutionary relationships and structural integrity but is limited to well-studied protein families with abundant alignment data, making it less applicable for generating novel proteins with unknown homologs. To address the limitations of MSA dependency, researchers developed sequence-based approaches such as ProteinRNN, which treats protein as sequential data without the need for MSAs^29^. While ProteinRNN captures short-term dependencies in amino acid sequences, it struggles with long-range interactions and global structural consistency, which are critical for protein functionality.

Inspired by advancements in natural language processing (NLP), researchers adapted pre-trained transformer models for protein sequence generation. ProtBERT (Elnaggar et al., 2019) applies the BERT architecture to learn complex contextual and dependency representations of amino acids sequences from large protein databases^30^. However, ProtBERT primarily focuses on understanding existing sequences rather than generating entirely novel proteins. It is highly effective for protein classification and function prediction, it does not explicitly generate new sequences. Building upon the success of ProtBERT, ProGen (Madani et al., 2023) employs a large-scale transformer model trained on 1 billion protein sequences to generate diverse and functional protein^18^. ProGen incorporates controllable generation by fine-tuning on curated datasets, allowing for property-specific protein design. However, it requires massive datasets and lacks structural validation, making it challenging to ensure generated sequences retain functionally important motifs.

To enhance sequence diversity while maintaining structural feasibility, researchers explored deep generative models such as Variational Autoencoders (VAEs) and Generative Adversarial Networks (GANs). ProteinVAE (Zhao et al., 2020) utilizes a VAE framework to encode protein sequences into a continuous latent space and generate novel sequences by latent space decoding^31^. While ProteinVAE effectively captures the underlying distribution of protein families, the method often produce blurry or functionally inconsistent sequences. To overcome the limitations of VAEs, ProteinGAN (Lee et al., 2021) employs a GAN-based model with adversarial training to enhance the quality of protein sequence using a generator-discriminator framework^20^. However, GANs are prone to mode collapse, where the model generates highly similar sequences rather than exploring diverse functional possibilities. Additionally, GAN models often suffer from training instability.

More recently, diffusion models have gained traction in protein generation. ProtDiff (Nguyen et al., 2022) employs a diffusion-based approach to generate highly diverse and structurally viable proteins by iteratively denoising protein sequences^21^. Diffusion models offer better training stability compared to VAEs and GANs, but they are computationally expensive and require careful fine-tuning to achieve optimal performance.

Recognizing the importance of structural constraints in protein design, researchers have developed structure-aware generative models. TaxDiff introduces a taxonomic-guided diffusion model for controllable protein sequence generation^32^. This methodology involves a denoising process where Gaussian noise is added and then incrementally reduced, guided by taxonomic data. TaxDiff generates structurally stable proteins within specific taxonomic groups, enhancing the controllability and precision of protein sequence generation. By incorporating taxonomic information, TaxDiff ensures that generated proteins are not only novel but also biologically relevant within their respective taxonomic contexts. However, the reliance on accurate taxonomic data limits its applicability to well-characterized taxonomic groups. ProteinMPNN generates sequences optimized for specific target folds, ensuring the resulting proteins adopt biologically feasible structures^33^.

Large-scale transformer-based protein language models have significantly advanced protein sequence analysis and design by leveraging vast datasets and sophisticated attention mechanisms. Inspired by large language models (LLMs) like GPT-3, ProtGPT is trained on extensive protein databases, allowing it to generate highly plausible and functional protein sequences with minimal supervision^19^. While it demonstrates strong generalization capabilities, the integration of LLMs in protein design remains in its early stages, requiring further refinement to improve sequence diversity, functional accuracy, and structural validation.

Another foundational model in this domain is Evolutionary Scale Modeling (ESM-1b), introduced by Rives et al. (2021)^34^. ESM-1b, a transformer-based model trained on billions of protein sequences, effectively captures evolutionary relationships and structural information. While originally designed for protein function prediction and structural inference, ESM-1b has also shown promise in sequence generation by leveraging its deep contextual understanding. More recently, ESMFold has extended these capabilities by integrating largescale protein language models for both structure prediction and sequence generation, offering a fully automated, end-to-end pipeline for protein design^35^. By leveraging transformer-based architectures, ESMFold bridges the gap between sequence generation and structural feasibility, enabling in silico protein engineering with higher accuracy. However, similar to other language models, it lacks direct optimization for functional constraints, making it less effective for designing highly specialized proteins like Cas9 and Cas12.

While prior generative models have made significant advances in protein design, they exhibit several limitations: 1) MSA-dependent models (e.g., MSA Transformer) require extensive alignment data, limiting their applicability to novel protein families. 2) Transformer-based models (e.g., ProGen, ProtGPT) focus on sequence generation but lack explicit structural validation. 3) GANs, VAEs, diffusion models struggle with mode collapse and loss of functional integrity in generated sequences. 4) Structure-aware models (e.g., ProteinMPNN, ESMFold) are primarily focused on fold prediction, not de novo sequence generation.

To address these limitations, we propose CasGen, a Transformer-based deep generative model for designing novel Cas protein variants. Our approach integrates: 1) classification-guided latent space regularization, ensuring clear separation between Cas and non-Cas proteins to enhance sequence specificity; 2) margin-based latent space regularization, preserving well-defined boundaries between Cas and non-Cas regions with space margins, thereby improving the quality of generated Cas proteins in learned regions of the latent space; 3) Bayesian optimization to efficiently explore the latent space and refine the generation of functionally relevant sequences; 4) comprehensive validation using AlphaFold2 and AlphaFold3, confirming that the generated Cas proteins retain essential structural and functional characteristics.

By combining deep generative modeling, latent space optimization, and structural validation, the proposed CasGen represents a significant advancement in AI-driven Cas protein design, expanding the genome-editing toolkit with novel, functional variants that can enhance genome engineering applications.

## 3 Results

As illustrated in Figure 1, our design pipeline employs a transformer-encoder-based model that encodes both Cas and non-Cas protein sequences into a latent space while simultaneously learning to reconstruct the original sequences. Bayesian optimization is then utilized to efficiently explore the latent space, identifying feature vectors to generate Cas-like proteins. Once classified as Cas, these latent feature vectors are decoded into amino acid sequences and subsequently evaluated using structural prediction tools such as AlphaFold2^26^ to assess foldability and functional plausibility. To ensure novelty, sequences exhibiting excessive similarity to known Cas proteins are filtered out via sequence alignment tools (e.g., BLAST), refining the selection to genuinely novel candidates.

**Figure 1.**
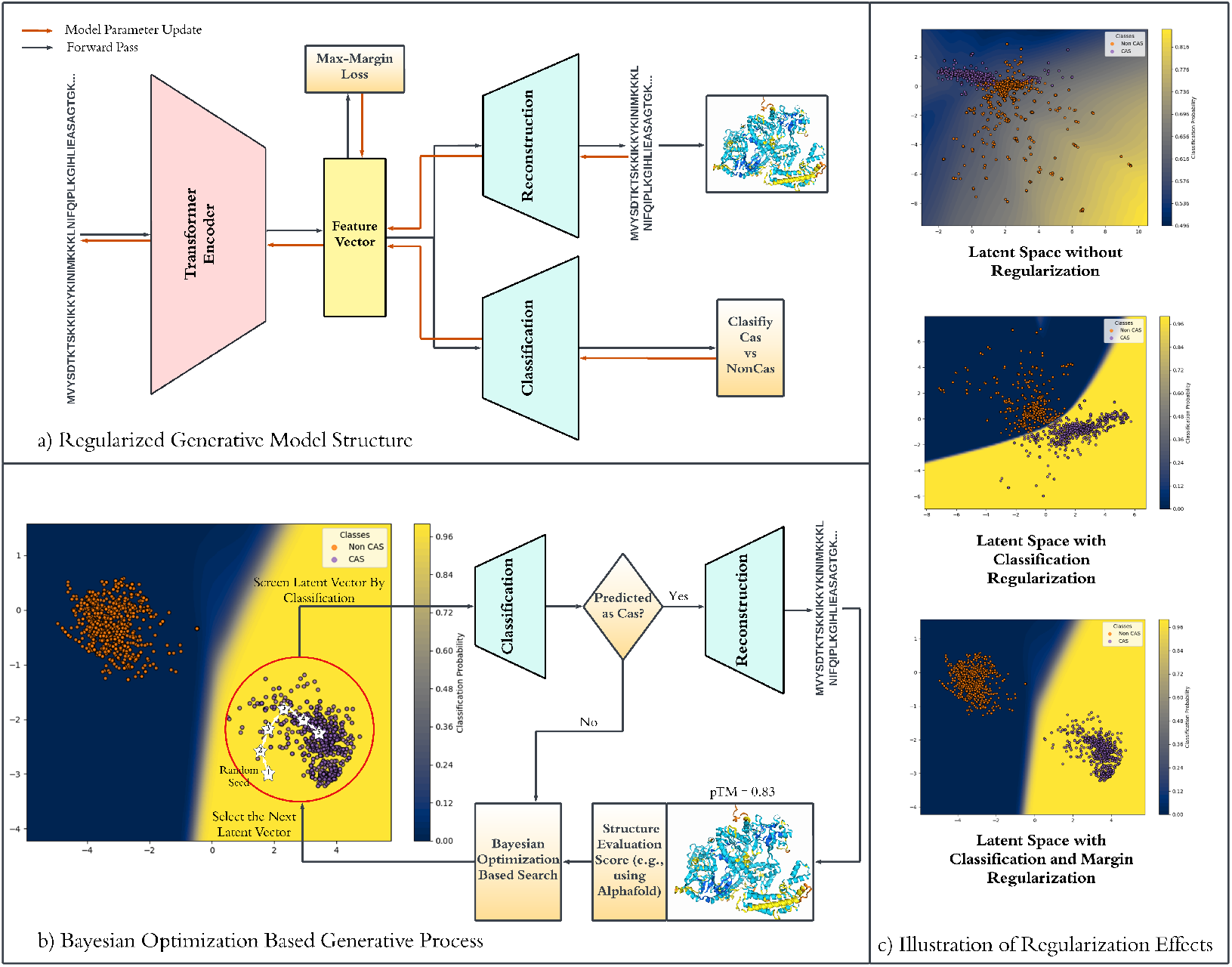
Overview of the Entire Model Procedure. a) Represent the Generative Transformer Model, b) illustrates the sampling technique used for sampling novel sequences, c) illustrates the effects of regularization of the latent space for No regularization, then regularization based on classification network and regularization based on classification network and margin loss

Through this iterative loop of classification, reconstruction, and structural validation, the final set of putative Cas proteins achieves an optimal balance between novelty and structural integrity, paving the way for further experimental validation.

### Sequence Alignment

The sequences generated by deep learning-based generative models were analyzed using an inhouse BLAST alignment pipeline against established protein databases, including NCI and Protein Data Bank (PDB). This sequence alignment step filters out sequences with high similarity to existing proteins, allowing for the selection of novel candidates for further structural and functional analysis.

### Structural Modeling and Analysis

The filtered sequences were used to generate three-dimensional structures utilizing AlphaFold2 (AF2), an AI-based structure prediction tool. To assess their structural similarity to known Cas9 or Cas12 proteins, the predicted structures were further analyzed using the FoldSeek^36^. This structural comparison helps identify regions of interest and evaluate the uniqueness of the designed proteins. The structural similarity was quantified using the TM-score, a metric for assessing topological resemblance between protein structures. The TM-scores ranged from 0.70 to 0.85, indicating a high degree of structural conservation with known structures, as values above 0.5 generally signify similar folds.

### Sequence Identity Analysis

To further assess the novelty of the generated proteins, sequence identity percentages were calculated relative to known Cas9 and Cas12 orthologs. The generated Cas9 variants (SpCas9, Nme1Cas9, and CjCas9) exhibited sequence identities ranging from 28% to 55%, while the Cas12 variants showed a maximum sequence identity of approximately 48%. These results confirm that the designed proteins are distinct in sequence while retaining key functional elements. This analysis was essential in verifying that the generative model produced novel Cas proteins rather than simply replicating known sequences.

### Structural Overlay and Comparison

Figure 2 illustrates the overlay of the designed Cas9 orthologs (SpCas9, Nme1Cas9, and CjCas9) with their respective known crystal structures. The structural alignment confirms that despite sequence variations, the overall structural integrity of the designed proteins closely resembles that of their natural counterparts. The high TM-scores (0.70-0.80) further validate that the generative model successfully produces structurally coherent and functionally plausible Cas protein variants, reinforcing the potential applicability of these designs in genome editing.

**Figure 2.**
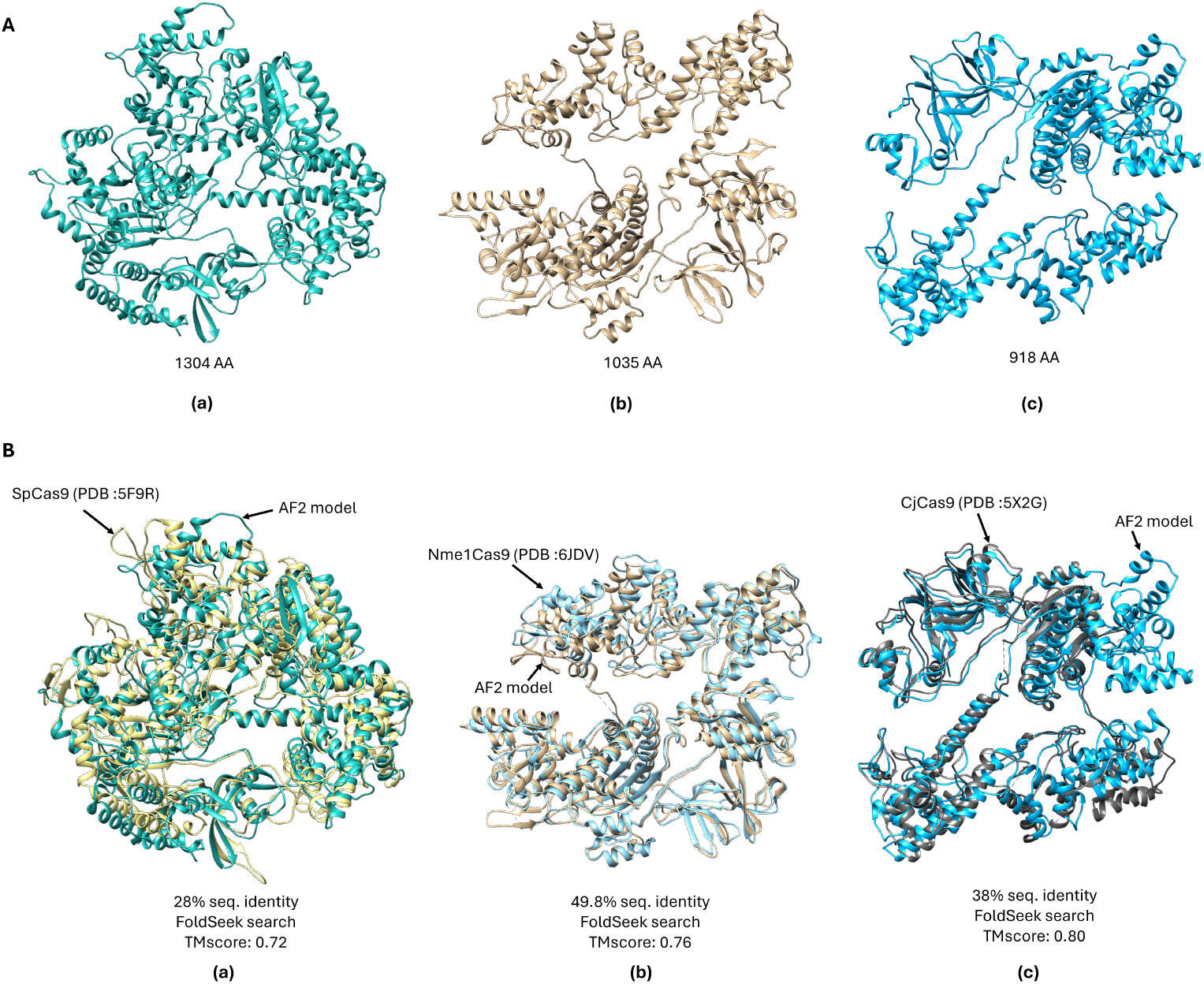
A). Alphafold2 (AF2) models of designed Cas9 orthologs (a) SpCas9 (b) Nme1Cas9 and (c) CjCas9 with their total amino acids (AA) sequence lengths B). Overlay of the designed Cas9 orthologs with known crystal structures from RCSB PDB.

### Tertiary Structure Prediction

To further evaluate the structural integrity of the engineered Cas9 and Cas12 proteins, their tertiary structures refined using AlphaFold3, incorporating guide RNA (g-RNA) and double-stranded DNA (ds-DNA) from their respective crystal structures. This additional modeling provided deeper insights into the interactions between the designed proteins and their nucleic acid substrates, as illustrated in Figure 3. The results demonstrate that our generative approach successfully produces novel Cas proteins capable of recognizing and interacting with g-RNA and ds-DNA targets in a manner consistent with their natural counterparts.

**Figure 3.**
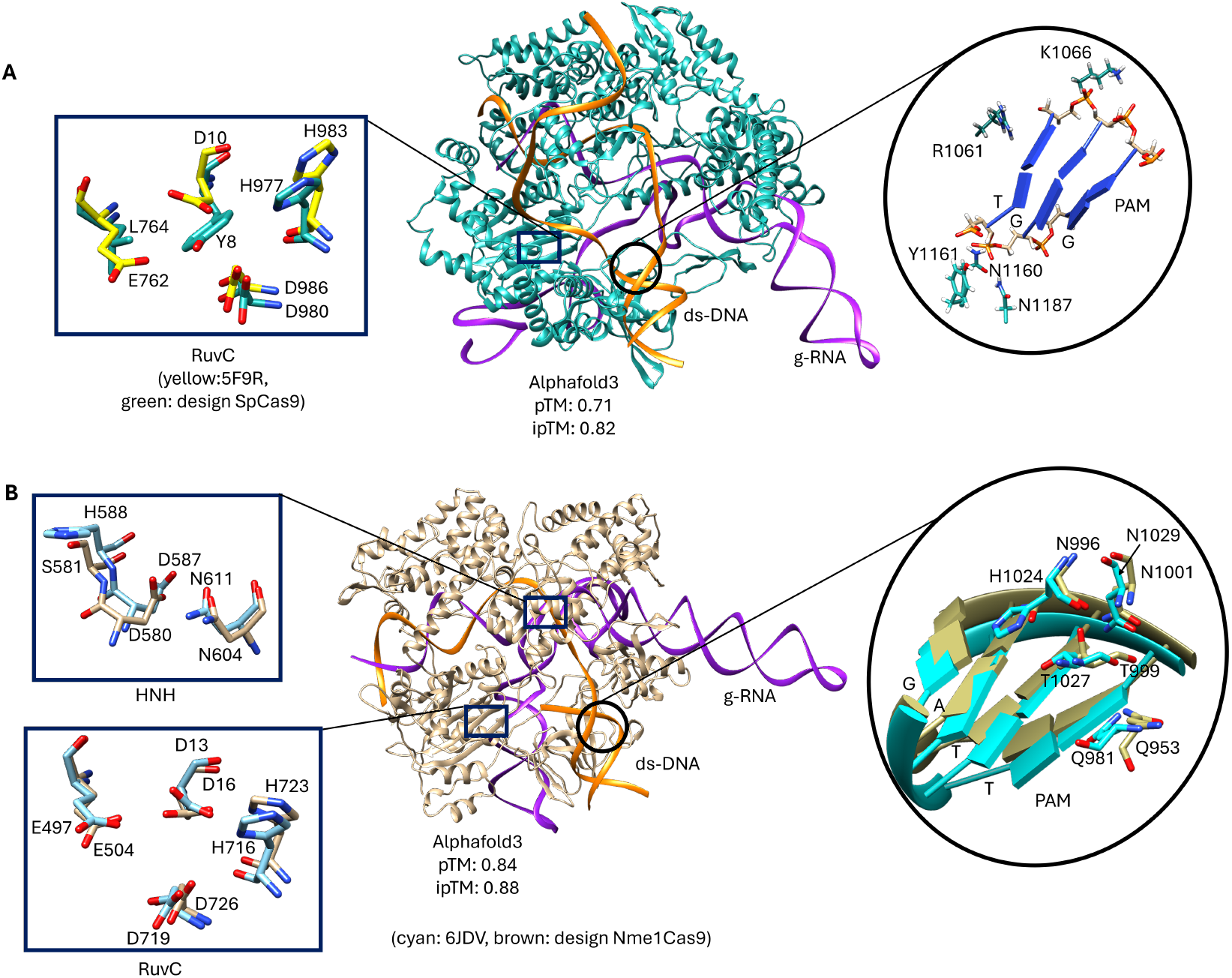
Alphafold3 models of tertiary structures (with corresponding ds-DNA and g-RNAs) of designed Cas9 orthologs A). SpCas9 B). Nme1Cas9 showing critical catalytic (HNH & RuvC) and functional (PAM) sites.

Notably, the tertiary structures of the designed SpCas9 and Nme1Cas9 orthologs exhibited a high degree of structural similarity to their respective crystal structures (PDB: 5F9R and 6DJV), with root-mean-square deviation (RMSD) values below 2.00 Å. In addition, the predicted template modeling (pTM) and interface predicted template modeling (ipTM) scores, predicted using AlphaFold3, ranging from 0.70 to 0.90, further confirming the structural accuracy of our designed Cas proteins. The strong structural resemblance in key functional domains, particularly in DNA binding and cleavage regions, suggests that these engineered orthologs retain their biological functions despite sequence variations.

Figure 3 illustrates the structural analysis of these designs, demonstrating that even orthologs with lower sequence identity exhibit a strong resemblance to natural Cas9 structures in key functional domains, including the RuvC nuclease, HNH nuclease, and protospacer adjacent motif (PAM)-interacting domains. For example, the designed Nme1Cas9, which has 49.8% sequence identity, retains key catalytic residues essential for HNH and RuvC nuclease domain formation as shown in Figure 3B. Furthermore, the PAM domain interaction, which involves a single residue change (H1024 to N996), remains functional integrity needed to initiate ds-DNA unwinding and reinforce the reliability of our design.

Conversely, the designed SpCas9, with a lower sequence identity of 28%, retains a functional RuvC nuclease domain with only two residue mutations (Figure 3A). In this model, the PAM interactions differ from the known SpCas9 structures, as many interactions are nonspecific, relying on the backbone of the PAM nucleotides. This observation highlights the uniqueness of our proposed design approach, demonstrating how sequence modifications can lead to alternative interaction strategies while maintaining structural feasibility.

Overall, the findings from this study highlight the effectiveness of using deep learning generative models with advanced structural prediction and analysis tools to design novel Cas proteins with functional and structural coherence. The validation against known crystal structures and subsequent structural analysis confirm that the generated proteins are both unique and maintain key functional characteristics. Furthermore, our CasGen framework successfully produces diverse Cas protein candidates that closely resemble naturally occurring variants in both sequence and structure, setting the stage for further experimental validation and potential applications in genome editing and protein engineering.

## 4 Discussion

In this study, we successfully developed a Cas protein generative model to design novel Cas9 and Cas12 protein sequences, aiming to expand the repertoire of genome-editing tools with enhanced or distinct functionalities. By leveraging the latent-space regularized generative transformer Model, we generated a diverse set of protein sequences that preserve essential functional characteristics while introducing novel sequence variations.

To ensure the novelty and structural viability of the generated proteins, we conducted comprehensive computational validation. Sequence alignment using in-house BLAST method ensured that the designed sequences were not redundant recreations of known proteins but instead novel entities with potential functional diversity. By filtering out sequences with high similarity to known proteins in established databases such as NCI and PDB, we prioritized designs that could contribute meaningfully to genome-editing advancements. Subsequent structural modeling with AlphaFold2 and AlphaFold3 revealed that the generated proteins retained significant structural integrity, as evidenced by high TM-scores (0.70–0.85) and low RMSD values (2.00 Å) relative to known crystal structures. These metrics confirm that the proposed CasGen approach effectively maintains key structural elements, ensuring that the generated Cas proteins remain functionally viable while permitting sequence-level diversity.

The structural overlays and tertiary structure predictions further demonstrated that our CasGen approach successfully produces proteins that not only mimic the structural folds of natural Cas proteins but also maintain critical interaction sites essential for DNA binding and cleavage. Our Cas design approach emphasizes the design of sequence variants that maintain essential domains while introducing mutations in regions less critical for core functions. The ability to preserve structural motifs critical for Cas9 and Cas12 activities while permitting diversity in other regions illustrates the robustness of our proposed generative process. These findings highlight the potential of integrating advanced machine learning techniques with state-of-the-art structural prediction tools in the field of protein engineering. Beyond preserving essential motifs, this framework provides a valuable tool for exploring novel protein functionalities that extend beyond naturally occurring sequences. By leveraging deep generative modeling in conjunction with advanced structural prediction tools, we enable a systematic and scalable approach to protein engineering, paving the way for customized genome-editing technologies with diverse applications in biomedicine, synthetic biology, and precision therapeutics.

While our study provides a robust computational framework for designing functional Cas protein variants, experimental validation remains a crucial next step. In vitro and in vivo assays are necessary to empirically assess the biological activity, genome-editing efficiency, and specificity of the generated Cas9 and Cas12 orthologs. Additionally, evaluating potential off-target effects will be critical for ensuring the safety and precision of these novel genome-editing tools. In future work, we will synthesize selected protein candidates and conducting in vitro and in vivo assays to evaluate their genome-editing capabilities. These experimental results will also serve as feedback for iteratively refining our generative model, incorporating empirical data to enhance prediction accuracy and optimize future protein designs. Integration of functional screening techniques with generative AI could lead to adaptive learning models, capable of dynamically improving protein generation based on real-world validation data.

In conclusion, our approach demonstrates the promising synergy between deep learning generative models and computational validation methods in advancing genome-editing technologies. This framework not only facilitates the discovery of novel genome-editing tools but also paves the way for the rational design of proteins with tailored functionalities for diverse biomedical applications. By integrating machine learning, structural modeling, and experimental validation, this study establish a scalable and iterative design pipeline for de novo Cas protein design. Our approach represents a significant step forward in the development of new genome-editing tools, offering new opportunities for precision medicine, synthetic biology, and next-generation therapeutic interventions. As experimental validation progresses, this methodology has the potential to transform the landscape of genome engineering, enabling the development of highly optimized, programmable Cas proteins with enhanced specificity, efficiency, and novel functionalities.

## 5 Methods

### 5.1 Dataset

To facilitate the design of novel Cas9 and Cas12 variants, we curated a comprehensive dataset comprising 3,021 Cas9 protein sequences, 287 Cas12 protein sequences, and 597 NonCas protein sequences from reputable biological databases. All Cas9 and Cas12 sequences were obtained from the InterPro database^37^, while Non-Cas proteins were sourced from diverse functional families, including DNA endonucleases, proteases, exonucleases, and helicases. This diverse dataset ensures a broad representation of enzymatic functions and structural features, which is essential for training robust classification and generative models.

To ensure the accuracy and biological relevance of the dataset, all Cas9 and Cas12 sequences underwent rigorous domain verification to confirm the presence of key functional domains. For Cas9 proteins, verification focused on the presence of RuvC and HNH nuclease domains, as well as the PAM-interacting domain, which are essential for DNA cleavage. For Cas12 proteins, domain validation included the RuvC nuclease domain and other functionally significant motifs specific to Cas12 orthologs. Sequences lacking these critical domains were excluded from our study, ensuring that only functionally relevant Cas proteins were incorporated into the dataset. This verification process was essential for generative model training, as it ensured that Cas-like proteins were accurately represented, improving generative protein design precision and performance.

The dataset exhibits a class imbalance, with a significantly higher number of Cas9 sequences (3,021) compared to Cas12 (287) and Non-Cas proteins (597). This imbalance could lead to model bias, where the classifier disproportionately favors the dominant Cas9 class while struggling to generalize across Cas12 and Non-Cas proteins. To mitigate this, we applied class-weighted loss functions that assigned higher penalties to misclassifications of Cas12 and Non-Cas sequences. Weights were computed based on inverse class frequencies, ensuring that the model learned to distinguish all three classes effectively.

This curated and validated dataset serves as the foundation for developing and training our Cas generative models. The diverse and high-quality of the dataset allow machine learning models to capture intricate sequence patterns and structural nuances, which are essential for accurate classification and the generation of functionally novel Cas variants. Furthermore, the broad representation of enzymatic functions within the dataset mitigates potential biases and enhances model generalization, and could facilitates the discovery of Cas9 and Cas12 variants with improved genome-editing capabilities.

### 5.2 Protein Sequence Tokenization

We conceptualize protein sequences as a “language,” drawing inspiration from advanced natural language processing (NLP) techniques to uncover intricate patterns embedded within protein data. Just as sentences in NLP are composed of words, each protein sequence can be represented as a string of “words,” where each word corresponds to an amino acid. To achieve this, we define a vocabulary of 20 symbols representing the 20 standard amino acids using IUPAC one-letter notation, along with four special symbols to denote beginning, end, unknown, and padding regions within a protein sequence. The specific numeric encoding assigned to each amino acid and special token is detailed in Table 1.

**Table 1.** Protein Tokenization Table.

| Token | Value | Token | Value | Token | Value |
| --- | --- | --- | --- | --- | --- |
| A | 3 | M | 13 | V | 20 |
| C | 4 | N | 14 | W | 21 |
| D | 5 | P | 15 | Y | 22 |
| E | 6 | Q | 16 | X | 23 |
| F | 7 | R | 17 | < start > | 1 |
| G | 8 | S | 18 | < pad > | 0 |
| H | 9 | T | 19 | < end > | 2 |
| I | 10 |  |  |  |  |
| K | 11 |  |  |  |  |
| L | 12 |  |  |  |  |

To ensure consistency across all protein sequences, we standardized sequence lengths to 1,600 tokens by padding shorter sequences with special tokens. This length was determined based on an analysis of our dataset, which revealed that all collected Cas sequences fall within this range. The sequence length standardization ensures that all sequences have uniform input dimensions, which is essential for efficient deep learning model training and evaluation, and handling of variable sequence lengths. Once tokenized, each protein sequence is transformed into a numerical representation using the predefined encoding scheme outlined in Table 1. This transformation is critical, as deep learning models require structured inputs to extract meaningful patterns, identify functional motifs, recognize conserved domains, and capture structural interactions essential for protein properties, such as genome-editing functionality.

### 5.3 Transformer-Based Protein Generative Model

Leveraging the remarkable success of Transformer models in natural language processing (NLP), we developed a specialized architecture for sequential protein language data analysis. Our approach integrates a Transformer encoder-based architecture with a 1D Convolutional Neural Network (CNN) decoder, paralleling the ReLSO (Regularized Latent Space Optimization) model^38^. This hybrid model capitalizes on the Transformer’s strength in deep sequence encoding and the CNN’s effectiveness in sequence reconstruction to facilitate protein generation tasks.

At the core of our model is the Transformer encoder, which processes tokenized amino acid sequences to extract rich, contextual representations. Each layer of the Transformer encoder comprises a multi-head self-attention mechanism and a fully connected feed-forward network, both enhanced with residual connections and layer normalization, enabling the model to simultaneously focus on different regions of the protein sequence, capturing intricate dependencies and long-range interactions between amino acids. To ensure training stability and mitigate overfitting, dropout layers are incorporated within each Transformer encoder block. Following the encoding process, the 1D CNN decoder reconstructs the original protein sequences from their latent representations. The decoder employs a series of convolutional operations to capture local sequence patterns, translating deep representations in latent space back into coherent amino acid sequences.

### 5.4 Latent Space Regularization

While transformer-based autoencoder frameworks excel at extracting high-level representations of protein sequences, they face significant challenges in maintaining a structured and well-separated latent space for the controlled generation of target proteins with desired properties. Without explicit regularization, the high-dimensional latent space becomes unstructured and entangled, Cas and non-Cas protein representations may highly overlap, making it difficult to differentiate between Cas and non-Cas proteins. The lack of organization and class separability in latent space leads to unreliable generative sampling, as new protein sequences drawn from an unregularized latent space may not retain functional properties or structural validity for desired target proteins. An unstructured latent space with poor class separability leads to unreliable generative sampling, where newly generated protein sequences may fail to retain essential functional properties or structural validity for the desired target proteins. These challenges are particularly critical when the model must generate novel Cas9 or Cas12 sequences and verify their classification by comparing them with seed sequences. If the latent space is not well-structured, generated sequences may drift toward non-Cas regions, leading to low-quality or functionally irrelevant outputs. Inspired by the Regularized Latent Space Optimization (ReLSO) model, which jointly trains an autoencoder and a prediction network to structure the latent space for improving the fitness scores of generated protein sequences, we adopt a similar architectural paradigm. However, in this study, we introduce classification-based and margin-based regularization to explicitly organize the latent space, ensuring a clear separation between Cas and non-Cas proteins and a well-defined Cas protein region to enhance novel Cas protein generation with improved robustness and design stability. Figure 4 illustrates the architecture of the proposed Latent Space Regularized Max-Margin Transformer Model (LSRMT), showcasing the interplay between the Transformer encoder, CNN decoder, classification network, and margin regularization module.

**Figure 4.**
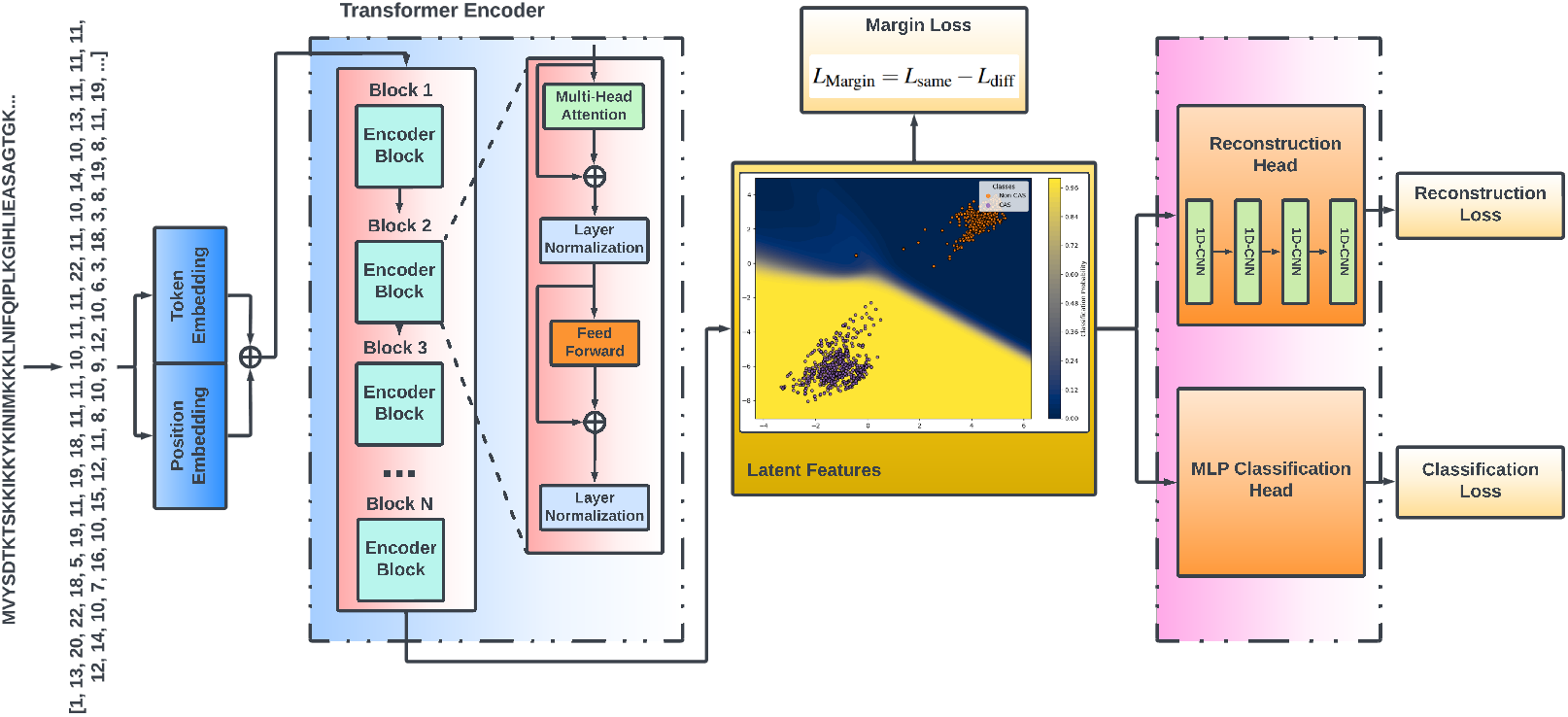
Regularized Generative Transformer Model (RGTM)

#### 5.4.1 Classification-Based Regularization

To ensure a well-structured latent space, we incorporate a multilayer perceptron (MLP)-based classifier network that regularizes latent representations of Cas and non-Cas proteins. By aligning sequence encoding with classification objectives, our method explicitly separates Cas and non-Cas regions within the latent space, improving class distinction and facilitating more precise target sequence discrimination. This approach enforces class-aware organization in latent space, ensuring that Cas proteins occupy distinct and well-defined regions in the latent space. By reducing overlap between classes, the model achieves a clear boundary for Cas protein generation.

We define the classification task as a binary classification problem with two classes:

- Class 1: Cas proteins (Cas9 or Cas12)
- Class 0: Non-Cas proteins

For a given protein *i*, let **X**_*i*_ ∈ ℝ^*d*^ denote its *d*-dimensional latent feature representation, which is computed by the Transformer encoder. The MLP classifier maps this latent representation to a logit vector **z**_*i*_ = [*z*_*i*1_, *z*_*i*0_] corresponding to Cas and non-Cas classes:

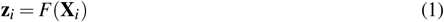

where *F*(·) represents the transformation applied by the MLP classifier network. Applying the softmax function to **z**_*i*_ converts the logits into class probabilities:

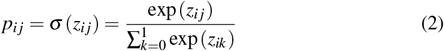

where *p*_*i j*_ represents the probability that sample *i* belongs to class *j* ∈ {0, 1}.

##### Cross-Entropy Loss for Classification

to train the classifier, the Cross-Entropy Loss is used to measure how well the predicted probabilities match the true class labels. Given a mini-batch of *n* protein sequences, the loss function is defined as:

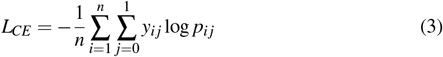

where *y*_*i j*_ is the ground truth label (binary indicator, 1 if sample *i* belongs to class *j*, otherwise 0), and *p*_*i j*_ is the predicted probability that sample *i* belongs to class *j*.

This classification-based regularization ensures that the latent space is well-structured for effectively differentiating between Cas and non-Cas proteins, while also enhancing the model’s ability to efficiently generate novel Cas proteins.

#### 5.4.2 Margin-Based Regularization

To enhance the structure of the latent space, we introduce a Max-Margin Latent Space Regularization strategy designed to enforce inter-class separability while promoting intra-class compactness. This approach ensures that embeddings of proteins from the same family remain tightly clustered while embeddings of different families are pushed farther apart. By structuring the latent space in this manner, the model improves its ability to generalize to unseen sequences while preserving essential functional properties across diverse protein families. To formalize this organization, we employ a Margin Loss Function, which directly influences the spatial configuration of embeddings by: 1) Minimizing intra-class distances, encouraging compact clusters of proteins within the same family; Maximizing inter-class distances, ensuring clear separation between Cas and non-Cas proteins. Max-Margin Regularization provides several key benefits. First, it enhances class separation by explicitly controlling the distance between Cas and non-Cas embeddings, ensuring precise differentiation for generative sampling. Second, it improves generalization by creating a more interpretable and functionally relevant latent space, enabling the model to better generalize to novel protein sequences. Finally, it reduces redundancy in generated sequences by promoting diversity in Cas9 and Cas12 variants, minimizing repetition while facilitating the discovery of novel functional proteins.

The margin Loss is computed using the following key steps:

- **Intra-Class Distance (Compactness)**: The intra-class distance measures the compactness of Cas and non-Cas embeddings. It is defined as the sum of the intra-class distances for Cas and non-Cas proteins:

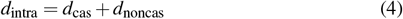

where the individual terms are computed as:

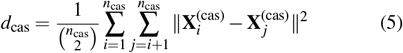

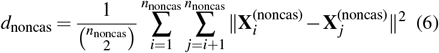

where *n*_cas_ and *n*_noncas_ are the number of samples in Cas and non-Cas classes, respectively.
- **Inter-Class Distance (Separation)**: The inter-class distance ensures clear separation between Cas and non-Cas embeddings by measuring the average squared pairwise distance between samples from different classes:

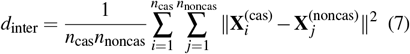

where 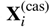 and 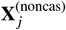 are the feature vectors of Cas and non-Cas proteins, respectively.
- **Total Weighted Margin Loss**: The final margin loss is computed by balancing intra-class compactness and inter-class separation through weighted contributions:

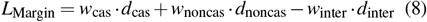

where *w*_cas_ and *w*_noncas_ control the compactness of Cas and non-Cas embeddings, respectively, while *w*_inter_ regulates the impact of inter-class separation. The Max-Margin Regularization improves the structure of the latent space, ensuring that sequences from the same family remain compact, while those from different families are well-separated. By structuring the latent space this way, the model generalizes better to unseen sequences while preserving essential functional properties across protein families.

### 5.5 Reconstruction Loss

During training, the model learns to reconstruct the input protein sequences by minimizing the Reconstruction Loss *L*_Recon_. We employ a CNN decoder, which takes latent representations from the Transformer encoder and generates output sequences that aim to match the original inputs. The fidelity of this reconstruction is measured using cross-entropy loss:

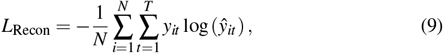

where: *N* is the number of samples, *T* is the sequence length, *y*_*it*_ represents the true token at position *t* in sequence *i*, 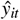 is the predicted probability of the token at the same position. This loss function ensures that the decoder learns to generate sequences that closely resemble the original protein sequences while preserving structural and functional integrity.

### 5.6 Total Loss Function

The total Loss combines the reconstruction loss, the crossentropy loss, and the margin loss defined as:

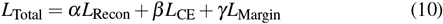

where *α, β* , and *γ* are weighting coefficients that control the relative contributions of each loss component: *α* (Reconstruction Loss Weight): Determines the importance of accurate sequence reconstruction, ensuring the decoder generates protein sequences that closely match the original inputs, *β* (Classification Loss Weight): Controls the emphasis on classification performance, influencing how well the model distinguishes between Cas and non-Cas proteins, and *γ* (Margin Loss Weight): Regulates the impact of latent space regularization, enforcing compactness within classes and separation between different protein families. By tuning these control parameters, the model can achieve a balanced optimization, ensuring high sequence reconstruction fidelity, classification accuracy, and well-structured latent space organization for reliable Cas protein generation.

### 5.7 Bayesian Optimization-Based Generative Process

The generation of novel, foldable protein sequences is accomplished through an iterative Bayesian optimization framework that integrates classification, sequence reconstruction, and structural evaluation. This approach systematically explores the latent space to identify latent feature vectors that correspond to functionally viable and structurally stable Cas proteins. The overall workflow is illustrated in Figure 1.

The generative process begins with the selection of an initial seed feature vector from the latent space encoded by the trained LSRMT model. This seed vector serves as a starting point for exploration. The classification network first evaluates the feature vector, predicting whether it corresponds to a Cas or Non-Cas protein: If classified as Cas, the feature vector is passed to the CNN-based reconstruction network to generate a candidate protein sequence. If classified as Non-Cas, the vector is returned to the Bayesian optimization module, which selects a new feature vector for exploration, as shown in Figure 1.

Once a feature vector is classified as Cas, the CNN-based reconstruction network decodes it into a corresponding amino acid sequence. This sequence then undergoes structural evaluation using predictive models such as AlphaFold^26^ or Boltz^39^ to assess its foldability and functional integrity. These models provide structural metrics, including the predicted TM-score (pTM) or TM-score, which indicate whether the generated sequence adopts a stable and biologically meaningful fold.

The Bayesian optimization module iteratively refines the selection of feature vectors by leveraging feedback from structure evaluation. The relationship between feature vectors and sequence quality (as determined by classification confidence and structure scores) is modeled using a probabilistic surrogate model, which is updated at each iteration. An acquisition function (e.g., Expected Improvement or Upper Confidence Bound) determines the next feature vector to explore, balancing exploration of new regions and exploitation of promising areas.

This Bayesian optimization-driven generative process offers several key advantages: 1) Targeted Sequence Generation: The integration of classification, reconstruction, and structural evaluation ensures that only Cas-classified proteins are reconstructed and assessed, reducing computational overhead. 2) Iterative Refinement: Bayesian optimization iteratively improves the selection of feature vectors, increasing the likelihood of generating structurally viable and functional Cas proteins. 3) Efficient Latent Space Exploration: The use of surrogate models and acquisition functions enables the model to efficiently navigate the latent space, significantly reducing the number of evaluations required compared to random search methods. 4) Structural Fidelity: Incorporating state-ofthe-art structure prediction models (such as AlphaFold and Boltz) ensures that only sequences with a high likelihood of proper folding are retained for further study. By iteratively refining the exploration of the latent space and ensuring that only structurally viable Cas proteins are selected, this approach streamlines the generative process and enhances the reliability of novel protein designs.

### 5.8 Validation Methods

To ensure the reliability and novelty of the protein sequences generated by our models, we employed a multi-stage validation pipeline that integrates sequence filtration, structural prediction, and comparative analysis.

The first step in the validation process involved filtering the generated protein sequences using a custom implementation of the Basic Local Alignment Search Tool (BLAST). This tool identifies regions of similarity by comparing sequences against established databases and assessing their statistical significance. Sequence identity percentages were evaluated, and only sequences with less than 50% identity to known proteins were selected as novel candidates for further analysis, ensuring that the generated proteins were not redundant copies of existing Cas9 or Cas12 variants.

To assess the structural integrity of the filtered sequences, their 3D structures were predicted using AlphaFold2 (AF2), an advanced AI-based protein structure prediction model. For each sequence, five structural models were generated, accompanied by per-residue confidence scores using the predicted Local Distance Difference Test (pLDDT), which evaluates the reliability and accuracy of the predicted structures. The predicted structures were further analyzed using FoldSeek, a structural search tool that aligns query structures to a database of known Cas9 and Cas12 proteins. FoldSeek provides quantitative similarity metrics, including the Template Modeling Score (TM-score), which measures structural similarity, and Root Mean Square Deviation (RMSD), which quantifies atomic positional differences between the predicted and reference structures.

Following this structural assessment, protein structures exhibiting high similarity to known Cas9 or Cas12 variants were superimposed onto their corresponding reference structures using Chimera and Yasara software^40,41^. These tools enabled precise structural alignment and visualization, allowing for a detailed comparison of conserved regions and overall structural congruence between the generated and known Cas proteins.

Finally, to refine the predicted tertiary structures and assess their interactions with nucleic acids, the engineered Cas9 and Cas12 proteins were modeled in complex with guide RNA (g-RNA) and double-stranded DNA (ds-DNA) using AlphaFold3. This advanced structure prediction tool integrates deep learning algorithms with molecular dynamics simulations to improve the accuracy of atomic interactions and conformational dynamics, resulting in high-resolution models of protein-DNA-RNA complexes. This refinement step ensures that the generated proteins not only adopt stable folds but also retain key functional attributes necessary for genome-editing applications.

By systematically integrating sequence filtering, structural modeling, and comparative validation, this multi-step approach ensures that the generated Cas proteins exhibit structural integrity, functional relevance, and a high degree of confidence in their folding and interaction properties.

## Code Availability

The CasGen model source code supporting this study is available at: https://github.com/shouyisxty/CasGen

## Acknowledgements

This work is supported by a grant from the National Institute of General Medical Sciences of the National Institutes of Health (R21GM144860).

## Author Contributions

Dr. Shouyi Wang and Dr. Jin Liu designed the study. Bharani Nammi implemented the methodology and conducted all computational experiments. Vindi Mahesha Jayasinghe Arachchige along with Maria Artiles, validated results. Sita Sirisha Madugula, Tyler Pham, and Charlene Norgan Radler curated and prepared the datasets. All authors contributed to the discussion, reviewed, and approved the final manuscript.

